# Frustraevo: A Web Server To Localize And Quantify The Conservation Of Local Energetic Frustration In Protein Families

**DOI:** 10.1101/2023.11.29.569273

**Authors:** R. Gonzalo Parra, Maria I. Freiberger, Miriam Poley-Gil, Miguel Fernandez-Martin, Leandro G. Radusky, Peter G. Wolynes, Diego U. Ferreiro, Alfonso Valencia

## Abstract

According to the Principle of Minimal Frustration, folded proteins can have only a minimal number of strong energetic conflicts in their native states. However, not all interactions are energetically optimized for folding but some remain in energetic conflict, i.e. they are highly frustrated. This remaining local energetic frustration has been shown to be statistically correlated with distinct functional aspects such as protein-protein interaction sites, allosterism and catalysis. Fuelled by the recent breakthroughs in efficient protein structure prediction that have made available good quality models for most proteins, we have developed a strategy to calculate local energetic frustration within large protein families and quantify its conservation over evolutionary time. Based on this evolutionary information we can identify how stability and functional constraints have appeared at the common ancestor of the family and have been maintained over the course of evolution. Here, we present FrustraEvo, a web server tool to calculate and quantify the conservation of local energetic frustration in protein families.

The webserver is freely available at URL: https://frustraevo.qb.fcen.uba.ar

## INTRODUCTION

Proteins are evolved systems that avoid strong energetic conflicts in their native states, thereby satisfying the Principle of Minimal Frustration ^1,2^. Although globular, folded proteins are globally minimally frustrated, local violations of this principle have evolved, opening up the possibility to encode the complex energy landscapes that enable active biological functions to emerge ^3^. Typical globular proteins have ∼10% of their residue-residue interactions in energetic conflict with their local surroundings; that is, they are highly frustrated^4^. This residual local energetic frustration (from now and on just frustration for simplicity) has been related to several functional aspects of proteins ^5^ such as protein-protein interactions ^4,6^, allosterism ^7^, catalysis ^8^ and order-disorder transitions ^9^, reinforcing the notion that protein physical stability and biological function sometimes conflict with each other.

Since AlphaFold2 was released, high quality structural models are available for a large majority of known proteins. Fueled by this availability, we have recently proposed a methodology to exploit the evolutionary information that is harbored within sets of homologous proteins, to identify energetic features of protein families that facilitate the biophysical understanding of a family and its individual members ^10^. Features that are common to a majority of family members can be interpreted as evolutionary conserved constraints. Based on the frustration states that are prevalent in those conserved regions, we can classify them into being mainly related to structural stability or functional requirements. Here we present FrustraEvo, a web server that applies information theory concepts to quantify frustration conservation in sets of proteins that belong to the same family. Through the lens of frustration, FrustraEvo’s results facilitate the interpretation of sequence variability within protein families and its impact in both structure stability and function.

## RESULTS

The algorithm to localize local energetic frustration was first introduced in 2007 ^4^ with an accompanying work where frustration patterns, and their consequences for the folding mechanisms of Im7, a protein with known 3D structure, were presented ^11^.

Im7 is an 87 amino acids single domain protein that inhibits the bacterial toxicity of its cognate protein, colicin E7. It has been shown that the residues that are located at the Im7 surface, on the interaction site with colicin E7, become highly frustrated when Im7 is alone. This frustration in turn has been shown to be correlated with an increased flexibility in that region ^2^. Moreover, this frustration is known to be linked to the appearance of an on-pathway folding intermediate state that is stabilized by non-native interactions ^11^. This high frustration in the native state of Im7 is released when the Im7-colicin E7 complex is formed. This is a clear example of the tradeoff that exists between stability and function in proteins ^3^. Residues that are used by evolution to perform specific molecular functions are often not favorable to local stability, making those proteins susceptible to detrimental phenomena such as aggregation or misfolding ^12^. Evolution has however found different strategies to minimize such risks, for example, through the use of chaperones. The Spy chaperone was shown to selectively recognise and interact with the highly frustrated residues within the interacting interface in Im7 ^13^. An unfrustrated state can subsequently be achieved by the interaction of Im7 with colicin E7, which releases it from the chaperone.

Structural information has been a limitation to study frustration on a large scale. The evolutionary context of proteins can be used to define regions that are constrained to change even when they are energetically detrimental, i.e. are highly frustrated. The recent advances in efficient structure prediction techniques ^14^, allow us to overcome such limitations and unblocked a whole new set of possibilities to perform high-throughput frustration analysis. In what follows we analyze the conservation of frustration levels within a set of Im7 homologs to better understand the evolutionary constraints of those proteins. In Fig. 1A the Multiple Sequence Frustration Alignment (MSFA) shows the distribution of both amino acid identities and frustration states for every aligned residue across the family members. The information content of the sequence (SeqIC) and frustration (FrustIC) are shown in the form of logos to visualize such conservation (Fig. 1B). Conservation of frustration levels is heterogeneous at different loci in the MSFA. Some residues show high variability both in sequence and in frustration e.g. position 17 that can be found as minimally frustrated, neutral or highly frustrated in different proteins of the family. This is reflected in the frustration logo (Fig. 1B) with no high FrustIC values for those positions. Some other regions can be variable in their amino acid identities but still be conserved in their frustration state, e.g. position 19 that although varying among the L, M, V or I amino acids stays in a minimally frustrated state. This is reflected with that position having a not so high SeqIC but a maximum FrustIC. Several positions composed of hydrophobic residues follow this pattern, highlighting their importance for the fold stability as being part of the hydrophobic core (green, tall bars in Fig. 1B). There are 3 residues that are highly frustrated and conserved in most members of the family (red, tall bars in Fig. 1B). Two of these residues are Y55 and Y56 that belong to the interacting region of Im7 with colicin E7 and Spy. In addition, W75 is also frustrated and conserved although it is not located at the protein-protein interaction site. W75, however, is reported to establish non-native interactions that are present at the intermediate state, possibly stabilizing it ^15^. If the intermediate state is an undesired consequence of the functional requirements associated to the binding residues used to bind colicin E7 why is there a residue that is detrimental to the local energetics in the native state that stabilizes it? Could it be possible that the intermediate itself has been positively selected? Notably, according to the MSFA, there is one protein in the dataset (WP199883497.1) that has that residue deleted. It would be of interest to perform MD simulations to assess whether this protein has such a folding intermediate state or not.

**Fig. 1.**
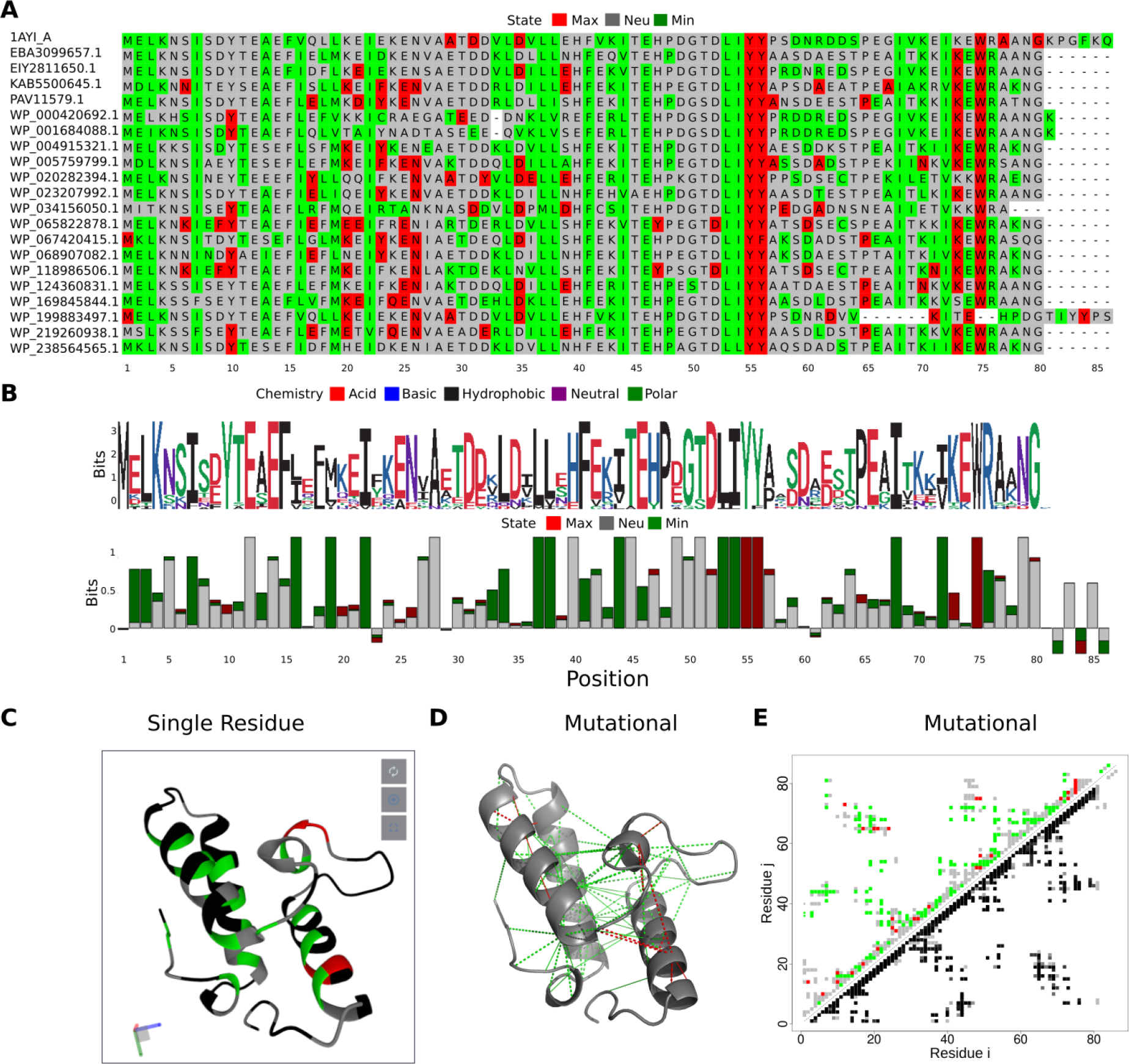
FrustraEvo outputs for the Im7 dataset. Different visualizations as produced by the FrustraEvo web server using PdbID=1AYI as the reference structure. **A**) Multiple Sequence Frustration Alignment (MSFA): Each residue in the MSA used as input is coloured according to its frustration state, mapped from the corresponding structure. **B**) Sequence and frustration logos: sequence and frustration conservation are visually represented as logos where SeqIC and FrustIC are displayed for each position. The individual contribution of each symbol to the total IC in each case is represented by the relative size of it to the size of the bars. **C**) Protein structure colored according to FrustIC based on SRFI as visualized on the web server. Residues with FrustIC >= 0.5 are coloured according to their most informative frustration value and residues with FrustIC < 0.5 are coloured in black. **D**) Protein structure of 1AYI with its contacts colored according to FrustIC values based on the mutational mode (only contacts with FrustIC >= 0.5 are shown) as it can be visualized in Pymol locally on the user’s computer. **E**) Contact maps: The FrustIC values, obtained from the contacts mode using the mutational FI are represented. In the upper diagonal matrix each point represents a contact between residues i and j where its color is assigned according to its most informative frustration state. In the lower diagonal matrix each point represents the same contacts as in the upper diagonal matrix but coloured in shadows of gray, representing the proportion of structures in the dataset that contain that contact between residues i and j.

### CONCLUDING REMARKS

The FrustraEvo web server implements a methodology that we have recently described to quantify the conservation of local energetic frustration in protein families ^10^. Residues whose frustration states are conserved in a majority of homologous proteins suggest the presence of selective pressure to maintain such a state. By exploiting knowledge of the evolutionary context of proteins, frustration conservation analysis allows researchers to identify structural or functional constraints in them at the level of single residues or pairwise interactions. Such constraints could be related to either primarily foldability and stability (mainly minimally frustrated patterns) or to functional reasons that conflict with folding (mainly highly frustrated patterns). FrustraEvo facilitates the interpretation of divergent signals in individual proteins as potential adaptations in their own evolutionary trajectories.

We are confident that frustration conservation analysis in protein families, as performed by FrustraEvo, will constitute a powerful tool that provides a biophysical interpretation that allows us to better understand the sequence-structure-function relationships in proteins.

## MATERIALS AND METHODS

### Input files

The protein structures need to be named using the same ID as their corresponding sequences within the MSA and the sequences between the structures and the corresponding entries in the MSA should match exactly. The protein structures can be derived from experiments or from structure prediction techniques like AlphaFold2 ^16^. Once both the MSA and the set of structures are uploaded into the server, users are asked to define a reference protein for the calculations. The reference will be used to define which columns in the MSA will be taken into account to make all downstream calculations in the pipeline. Only columns in the MSA that are not gaps in the reference protein are considered. More details can be found in ^10^. Finally, users can provide their email address to be notified when the job is completed. A job Id is also provided to retrieve the results. The web server is free and open to all and there is no login requirement.

### Server Calculations

As a first step FrustraEvo uses the R package FrustratometeR ^17^ to calculate frustration for all the uploaded structures. In a conceptual way frustration measures how well optimized for folding the energy of a residue or a given residue-residue interaction is in comparison to randomized structural decoys. The frustration index (FI), by residue or by contact, is calculated using a Z-score that compares the native energy according to the AWSEM energy function ^18^ with the distribution of the energies of 2000 decoys. The FI value can be classified into, 1) highly frustrated, if most decoys have a lower energy, 2) minimally frustrated, if most decoy possibilities have a higher energy or 3) neutral in an intermediate case scenario. According to the way the decoys are constructed we can obtain 3 different Frustration Indices (FIs). The mutational FI indicates how the frustration changes as a function of the amino acid identities for a given pair of positions. The configurational FI frustration measures how frustration changes not only relative to amino acid identities but also to the change of conformational state and solvent exposure. Although correlated, the mutational FI is somewhat more useful for identifying active or ligand binding sites, while the configurational FI can be employed to analyze protein-protein interactions and conformational changes. The Single Residue Frustration Index (SRFI) aggregates the energetic description of all interactions established by a given residue when mutating its identity. More details on how to calculate each FI can be found in ^4,19^.

Once frustration is calculated for the structures, FrustraEvo processes the MSA by keeping only those positions where the reference protein does not contain a gap. Frustration values are mapped from the structures to each residue from each protein contained in the MSA to generate a Multiple Sequence Frustration Alignment (MSFA) as shown in **Fig 1A**. To calculate frustration conservation at the single residue level, for each column in the MSFA, FrustraEvo calculates the Information Content (IC) for both sequence (SeqIC) and frustration (FrustIC). SeqIC is calculated from the MSA using the *ggseqlogo* R package ^20^. FrustIC is calculated from the distribution of frustration states within the MSFA using the following formula for each column:

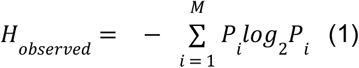

where *P*_*i*_ is the probability that the system is in frustration state i. The probabilities are normalized such as 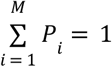, where M is the number of possible frustration states. (i.e. minimally, neutral or highly frustrated states, therefore, M=3). To take into account background probabilities, the information content is calculated as *FrustIC = H*_*max*_ *-H*_*observed*_ We used the following frequencies for the different frustration states as reported by Ferreiro et al ^4^ as background frequencies to calculate *H*_*max*_: minimally frustrated 0.4, highly frustrated 0.1 and neutral 0.5.

For the pairwise contacts mode, the reference structure is used to define the contacts to be evaluated. FrustraEvo calculates the frequency of having a contact between columns i, j in the MSA, on each structure in the dataset. Where i, j ∈ [1, N], with N being the number of columns in the ungapped MSA. FrustraEvo calculates, for each possible contact, the information content contributions from each frustration state. The FrustIC of a given contact will be calculated as the sum of the individual contributions from each frustration state. The background frequencies are the same as for the single residue mode.

### Outputs and Visualization

Results can be retrieved using either the JobID or the link that is provided at the time of data submission (which are also sent via email, if provided). In the job results page different visualizations are provided.

An MSFA is shown where SRFI results for all proteins can be visualized on top of the MSA **(Fig1A)**. This representation facilitates observing the different variability patterns among the proteins contained in the dataset to better understand the family features. The sequence and frustration logos summarizing the IC calculations as explained in methods are shown in Fig 1B where, based on the frustration states that are represented at each column in the MSFA, the sequence (SeqIC) or frustration (FrustIC) conservation values are shown. Mutational and Configurational FI conservation results are visualized in the form of contact maps (Fig1C shows the results for the mutational FI) where in the upper diagonal part of the matrix, each contact in the reference structure is represented as a point coloured according to its most informative frustration state. An important aspect to consider is that not all contacts are present in all structures. Therefore, in the lower diagonal matrix, the relative frequency with which the contact is present within the set of processed structures is shown in grayscale. Finally, FrustIC values are mapped into the reference structure and displayed to be interactively inspected in a Mol* session, based on the SRFI (Fig1D) as well as in the pairwise-contact FIs (Fig1E shows results for the mutational FI).

Plots for the frustration and sequence logos and contact maps are made using the *ggplot2* R package. Results can be downloaded by following the link at the bottom of the job page. Results are distributed as a zip file that contains the individual frustration calculations for all structures in the submitted dataset, raw tables summarizing the FrustIC as well as many other statistics based on the 3 different FIs, all the figures that are displayed in the server job page, Pymol scripts to interactively visualize FrustIC results on the different structures and a README file that explains the directory structure of the entire results files as well as how to interpret the raw tables that summarize the FrustIC results.

### Im7 Dataset

We used blastp within the NCBI blast web server to find Im7 homologous proteins ^21^, as a sequence query we used the PDB ID: 1AYI (chain A). We used default parameters and we searched against the non-redundant database of protein sequences. We took the 100 first hits from blastp and applied CD-Hit at 90% to reduce sequence redundancy. We obtained a total of 21 Im7 homologous sequences (Supplementary Table 1), from which only 1 had an available experimental structure deposited in PDB. Structures for the 20 sequences that do not have their structure solved were predicted using AlphaFold2.

## ACKNOWLEDGEMENTS

The authors would like to thank Miguel Romero, Camila Pontes and Victoria Ruiz-Serra for their useful comments and suggestions at different stages of the project. RGP holds a fellowship from Grant IHMC22/00007 funded by the Instituto de Salud Carlos III (ISCIII) and by the European Union NextGenerationEU/PRTR. DUF and MIF are supported by the Consejo de Investigaciones Cientificas y Tecnicas (CONICET). PGW is supported by the Center for Theoretical Biological Physics sponsored by the NSF grant PHY-2019745 and the D. R. Bullard-Welch Chair at the Rice University Grant C-0016.

